# *In Vitro* Characterization of Extracellular Vesicles from the Medicinal Plant *Centella asiatica* for Aesthetic Applications

**DOI:** 10.1101/2024.12.03.624435

**Authors:** Tsong-Min Chang, Chung-Chin Wu, Huey-Chun Huang, Shr-Shiuan Wang, Ching-Hua Chuang, Pei-Lun Kao, Wei-Hsuan Tang, Luke Tzu-Chi Liu, Wei-Yin Qiu, Ivona Percec, Charles Chen, Tsun-Yung Kuo

**Affiliations:** Department of Applied Cosmetology, HungKuang University, Taichung City, Taiwan; Schweitzer Biotech Company, Taipei City, Taiwan; Department of Medical Laboratory Science and Biotechnology, China Medical University, Taichung City, Taiwan; Department of Clinical Application, Center for iPS Cell Research and Application (CiRA), Kyoto University, Kyoto, Japan; Division of Plastic Surgery, Department of Surgery, University of Pennsylvania, Philadelphia, PA, USA; College of Science and Technology, Temple University, Philadelphia, PA, USA

**Keywords:** Centella asiatica, skin care, extracellular vesicles, antioxidants, cosmetics, anti-inflammatory, skin whitening, UV damage, photoaging

## Abstract

**Background:** *Centella asiatica* has long been used as a medicinal herb in traditional Asian medicine. Its wound healing, skin improvement, and neuroprotective properties have been widely studied. Extracellular vesicles (EVs) are secreted by cells and contain bioactive components with therapeutic properties.

**Objectives:** This study aims to characterize EVs isolated from *C. asiatica* tissue culture and investigate their therapeutic properties using *in vitro* assays and a UVB-induced damage mouse model.

**Methods:** EVs were isolated from *C. asiatica* tissue culture and characterized by nanoparticle tracking analysis (NTA) and scanning electron microscopy (SEM). Cytotoxicity, anti-oxidation, anti-melanin, and anti-inflammation of the EVs were evaluated by MTT assay, tyrosinase assay and RT-qPCR in chemical or *in vitro* assays. A UVB-induced photodamage mouse model was established to assess the anti-inflammation effect of EVs *in vivo*. Gels with or without EVs were applied to the damaged site and skin appearance was observed daily and skin histopathology was analyzed on day 7 by H&E and immunohistochemical staining.

**Results:** *C. asiatica* EVs were found to contain high levels of polyphenols and mitigate hydrogen peroxide-induced intracellular ROS. The EVs were further able to reduce intracellular melanin content and tyrosinase activity, and exhibited anti-inflammatory effects by downregulating the expression of pro-inflammatory genes COX2 as well as nitric oxide production. In mice with UVB-induced skin damage, daily application of *C. asiatica* EV gel reduced skin epidermis thickness and inflammation compared to UVB-only or blank gel at seven days after UV irradiation.

**Conclusions:** The beneficial effects of *C. asiatica* EVs on skin quality warrant further studies as promising agents in skin care applications.

## Introduction

*Centella asiatica*, also commonly known as Gotu Kola, is a traditional medicinal herb widely used across Asia [1]. Its broad spectrum of therapeutic properties is attributed to a diverse phytochemical profile, including triterpenoids, flavonoids, and polyphenols, which contribute to the plant’s anti-inflammatory, antioxidant, and wound-healing properties, as well as its neuroprotective and cognitive-enhancing effects in animal studies [2–4]. Several key triterpenes, including madecassoside, madecassic acid, asiaticoside, and asiatic acid, are well-studied compounds that play primary roles in the above therapeutic effects [4, 5]. Due to its wide-ranging therapeutic benefits, *C. asiatica* has garnered interest in modern pharmacology for potential applications as a natural treatment in chronic inflammation, neurodegenerative diseases, and skin improvement.

Extracellular vesicles (EVs) are membrane-bound particles of various sizes and compositions released by cells into the extracellular environment, serving as carriers of bioactive molecules such as proteins, lipids, and nucleic acids [6]. EVs are classified based on size, origin, and markers: exosomes (30–150 nm), which originate from the multivesicular bodies in the endosomal pathway; microvesicles (100–1000 nm), which bud directly from the plasma membrane; apoptotic bodies (1000-5000 nm), which are formed from membrane blebbing in apoptotic cells [6, 7]. These vesicles play essential roles in intercellular communication, modulating physiological and pathological processes by delivering molecular signals to recipient cells [6]. Due to their stability and biocompatibility, EVs have been explored for applications ranging from drug delivery and diagnostics to regenerative medicine, as well as skin therapies via non-invasive topical applications [7, 8].

Plant-derived EVs share similarities with mammalian EVs. For example, exosome-like particles in plants also form from multivesicular bodies, akin to processes in mammalian cells [9]. Recently, plant-derived EVs have attracted considerable attention as an alternative to mammalian EVs due to their comparable therapeutic potential and additional advantages in safety, scalability, cost-effectiveness, and ethical sourcing [9, 10]. Plant-derived RNA, particularly microRNAs (miRNAs), can be found in EVs and play a significant role by influencing gene expression in recipient cells, particularly via cross-kingdom communication, where they can modulate the expression of target genes in other organisms, including humans [11]. Recent studies have shown that exosome-like EVs from plants, including *C. asiatica*, promote skin improvement and wound healing more effectively than conventional plant extracts [12]. For instance, *C. asiatica* exosome-like EVs can upregulate the COLA1 collagen gene and downregulate the MMP1 matrix metalloproteinase gene, and inhibit melanogenesis genes [13]. Furthermore, *C. asiatica* exosome-like EVs can be absorbed into human live cancer cell culture within 24 hours and exhibit anticancer effects by promoting apoptosis of cancerous cells [14].

Many medicinal plants, including *C. asiatica*, are subject to overharvesting, which threatens biodiversity and ecosystem health. By using tissue culture techniques, EVs can be obtained without putting strain on ecological resources, supporting conservation efforts and ethical sourcing while meeting production demand [15]. In light of the therapeutic potential of *C. asiatica* and the unique properties of EVs as skin-penetrating biomolecular vehicles, this study aims to investigate the *in vitro* properties of EVs isolated from *C. asiatica* tissue culture as well as the effects of *C. asiatica* EV-based gel on UV-induced skin damage in a mouse model.

## Materials and Methods

### Isolation and characterization of *Centella asiatica* extracellular vesicles

*C. asiatica* EVs were extracted and isolated by a multi-step filtration and centrifugation. Briefly, tissue culture *C. asiatica* was homogenized, coarsely filtered to remove large debris, and centrifuged. Pellets were collected and filtered through a 0.45 μm filter, extracted and finally filtered through a 0.45/0.22 μm membrane filter.

The size and morphology of the *C. asiatica* EVs were analyzed with nanoparticle tracking analysis (NTA) and transmission electron microscopy (TEM). The NTA was performed with ZetaView® system (Particle Metrix, Germany). A sample for TEM was prepared by dropping 10 μL of the specimen onto a copper grid, followed by rest for 2 minutes and blotted dry with filter paper. Ten μL of uranyl acetate was then applied to the grid and excess liquid was again dried with filter paper, and the process was again repeated. Afterward, the grid was air-dried and subjected to the TEM procedure.

### Cell culture

The B16F10 cell line (mouse melanoma cells, ATCC CRL6475, BCRC60031) was obtained from the Bioresource Collection and Research Center (BCRC), Taiwan. The cells were maintained in Dulbecco’s Modified Eagle Medium (DMEM) (Hyclone, Logan, UT) supplemented with 10% fetal bovine serum (FBS) and 1% antibiotics. Human keratinocyte line HaCaT (AddexBio T0020001) was grown in high-glucose DMEM supplemented with 10% FBS, 4 mM/L-glutamine, and 1% antibiotic-antimycotic solution. Human foreskin fibroblast line Hs68 (BCRC 60038) was grown in high-glucose DMEM supplemented with 10% FBS, 4 mM/L-glutamine, and 1% antibiotic-antimycotic solution. Human keratinocytes line Detroit 551 (BCRC 60118) was grown in Minimum Essential Medium (MEM) supplemented with 10% FBS, 0.1 mM non-essential amino acids, and 1.0 mM sodium pyruvate. All cells were maintained at 37°C in a humidified atmosphere containing 5% CO₂.

### Cell viability assay

The 3-(4,5-dimethylthiazol-2-yl)-2,5 diphenyltetrazolium bromide (MTT) method was performed for B16F10 cells. Briefly, the cells were exposed to various concentrations of *C. asiatica* EVs for 24 hours, and the MTT solution was then added to the wells. The insoluble derivative of MTT produced by intracellular dehydrogenase was solubilized with an ethanol-DMSO (1:1) mixture. The absorbance of the wells at 570 nm was read using a microplate reader. For HaCaT and Detroit 551 cells, 1/10 volume of alamarBlue® (resazurin) reagent was added directly after 24 hours of EV treatment and incubated for 4 hours at 37°C. The absorbance was then read at 570 nm with 600 nm as a reference. In both assays, cell viability was calculated as the ratio of absorbance of the treated cells to control.

### Thiazoline-6-sulfonic acid (ABTS) free radical scavenging assay

The ABTS free radical assays were conducted as previously outlined [16]. A 7 mM stock solution of ABTS was reacted with 2.45 mM potassium persulfate, and the mixture was left to stand in the dark for at least 6 hours before use. The absorbance at 734 nm was measured immediately after mixing various concentrations of *C. asiatica* EVs (4×10^6^, 1×10^7^, and 2×10^7^ particles/mL) with 1 mL of ABTS solution to detect changes within 10 minutes. Vitamin C (0.9 mg/mL) and BHA (0.9 mg/mL) were used as positive controls.

### Intracellular reactive oxygen species (ROS) assay

The intracellular ROS assay was performed using an established method [17]. B16F10 cells were cultured in 24-well plates and treated with various concentrations of *C. asiatica* EVs (1×10^9^, 2.5×10^9^, and 5×10^9^ particles/mL), Trolox^®^ (0.5 mg/mL, positive control), or none (negative control) for 24 hours. The cells were then incubated with 24 mM H_2_O_2_ for 30 minutes to induce oxidative stress. After incubation, 2’, 7’-dichloro-fluorescein diacetate (DCFH-DA) was added to the cells and cultured for 30 minutes. After treatment, the cells were washed with PBS and trypsinized with trypsin/EDTA. The fluorescence intensities of DCF were measured at an excitation wavelength of 504 nm and an emission wavelength of 524 nm using a Fluoroskan Ascent fluorescent reader (Thermo Scientific, Vantaa, Finland).

### Total polyphenol content assay

Different concentrations of *C. asiatica* EVs (2×10^7^, 4×10^7^, 5×10^7^ particles/mL) and gallic acid standard were mixed with Folin-Ciocalteu reagent and incubated for 5 minutes. Sodium carbonate solution was added and incubated for 30 minutes. The absorbance at 765 nm was measured using a spectrophotometer and the gallic acid equivalent was calculated with the gallic acid standard curve [18].

### Metal chelating assay

The metal chelating activity of the extracellular vesicles was performed with potassium ferricyanide (K_3_[Fe(CN)_6_]) assay [19]. Different concentrations of *C. asiatica* EVs (2×10^6^, 5×10^6^, 1×10^7^ particles/mL) were mixed with 0.2 mM PBS and 1% (W/V) K_3_[Fe(CN)_6_], with 2 mg/mL EDTA as the positive control and deionized water as the blank control. The mixtures were heated at 50°C for 20 minutes. After cooling to room temperature, 10% (W/V) TCA was added, and the mixtures were centrifuged at 12,000 rpm and 4°C for 5 minutes. The supernatants were transferred to a 96-well plate, followed by the addition of deionized water and 1% (W/V) FeCl₃. The plates were incubated in the dark for 30 minutes. Absorbance was measured at OD 700 nm to assess metal chelating activity.

### Intracellular melanin content assay

The intracellular melanin content was measured as described previously [20]. B16F10 cells were treated with α-Melanocyte-stimulating hormone (α-MSH, 100 nM) for 24 h to induce melanin production. Following this, the cells were treated with different concentrations of *C. asiatica* EVs (1×10^9^, 2.5×10^9^, and 5×10^9^ particles/mL), or the positive control, arbutin (0.54 mg/mL), for an additional 24 hours. Post-treatment, cell pellets containing a known number of cells were solubilized in 1 N NaOH at 60°C for 60 minutes. The melanin content was then assayed at 405 nm.

### Intracellular tyrosinase activity assay

The intracellular tyrosinase activity was determined as described in previous research [21]. B16F10 cells were treated with α-Melanocyte-stimulating hormone (α-MSH, 100 nM) for 24 hours to induce melanin production. Following this, the cells were treated with different concentrations of *C. asiatica* EVs (1×10^9^, 2.5×10^9^, and 5×10^9^ particles/mL), or the positive control, arbutin (0.54 mg/mL), for an additional 24 hours. After treatments, the cell extracts (100 μL) were mixed with freshly prepared L-DOPA solution (0.1% in PBS) and incubated at 37°C. The absorbance at 490 nm was then measured to assess the tyrosinase activity.

### Type I procollagen (PIP) ELISA assay

Hs68 cells were seeded at 1×10^5^cells per well in 12 well plates and were cultured in 10% FBS DMEM medium at 37°C with 5% CO_2_ for 24 hours. After three washes with PBS, several different concentrations (1×10^7^, 1×10^8^, 1×10^9^ particles/mL) of *C. asiatica* EVs and 5 ng/mL TGF-β1 (AbCam Cat. #ab50036, as positive control) were added and cultured for 24 hours. The level of type I procollagen in cell culture supernatants was determined using a commercial PIP EIA Kit (TAKARA Cat. # MK101) according to the manufacturer’s instruction (2 step procedure).

### mRNA expression of aquaporin-3 (AQP3) and filaggrin (FLG) genes

HaCaT cells were dispensed into a 96-well plate and cultured for 24 hours under cell culture conditions. Various concentrations of *C. asiatica* EVs and negative control (purified water only) were added to the cells and cultured for 24 hours. Real-time PCR was performed to confirm the expression of aquaporin-3 (AQP3) and filaggrin (FLG) at the gene level, and the test sequence is as follows. For RNA isolation and cDNA synthesis, SuperPrep TM cell lysis & RT Kit for qPCR (TOYOBO, Cat. SCQ-101) was used. To compare and analyze gene expression, the cDNA synthesized above was used as a template, and Real-time PCR analysis was performed using Thunderbird TM SYBR qPCR Mix (TOYOBO, Cat. QPS-201). The gene expression levels of the sample were quantified by normalization with GAPDH using the 2^-△△Ct^ method.

### mRNA expression of pro-inflammatory genes after UVB induction

HaCaT cells (1×10^4^/well for 96-well) or Hs68 cells (5×10^4^/well for 24-well) were cultured for 24 hours and exposed to UVB at 90mJ dose. Various concentrations of *C. asiatica* EVs and negative control (purified water only) were added to the cells and cultured for 4 hours. Real-time PCR was performed to confirm the expression of cyclooxygenase-2 (COX) gene as above and the gene expression levels were quantified by normalization with GAPDH using the 2^-△△Ct^ method.

### Measurement of nitric oxide production

RAW 264.7 macrophage cells were seeded in 10% FBS DMEM medium in 24-well plates at a density of 2 × 10^5^ cells/well and were cultivated at 37°C with 5% CO_2_ for 24 hours. Removing the original medium, the cells were treated with 1×10^7^, 1×10^8^, 5×10^8^, 1×10^9^, 2×10^9^, or 4×10^9^ particles/mL of *C. asiatica* EVs or gallic acid (10 and 20 μg/mL, Sigma-Aldrich Cat. #G7384) in serum-free DMEM medium and incubated for 4 hours. After the pre-incubation, the cells were stimulated with 100 ng/mL LPS and incubated for 20 hours. The culture supernatants were harvested and centrifuged at 7,000 × g for 3 minutes. The samples were mixed with an equal volume (100 μL) of Griess reagent (Sigma-Aldrich Cat. #G4410) and the absorbance was measured at 540 nm after reacting for 10 minutes.

### Animal study

Twelve male ICR mice were obtained from BioLASCO Taiwan Co. Ltd. (Taipei City, Taiwan). The mice were 7 weeks old and weighed from 31 to 33 g at the start of the study. The mice were randomly assigned to four groups with three mice in each of the group as in Table 1.

**Table 1.** Group assignment for the animal study.

| Groups | Description | # of mice |
| --- | --- | --- |
| UVB | UVB exposure only | 3 |
| Control | UVB exposure + gel without active ingredients | 3 |
| Treatment 1 | UVB exposure + gel with purified *C. asiatica* EVs | 3 |
| Treatment 2 | UVB exposure + gel with purified *C. asiatica* EVs + TECA (Titrated extract of *C. asiatica*) | 3 |

To remove hair for test gel administration, mice were anesthetized with Isoflurane (3% – 4%), and back hair was shaved and further removed using hair removal cream. On Day 0, mice were exposed to 120 mJ/cm² UVB for 5 minutes to induce skin damage. From Day 0, after UVB exposure, the test gel was applied twice daily at a volume of 180–200 μL on the mice’s back according to the respective group assignments. Photographs of the hair removal area were taken before UVB exposure on Day 0 and daily from Day 1 to Day 7 before applying the gel to observe skin repair. Body weight of the animals were measured on Days 0, 1, 4, and 7. Animals were sacrificed on Day 7 for histopathological analysis.

### Animal tissue histopathological analysis

On the day of sacrifice, skin tissue samples were taken from back and fixed in 10% neutral formalin for at least 24 hours, embedded, and sectioned. The long axis of the skin tissue section was defined as the observation direction. After gradient alcohol dehydration, xylene permeation, and paraffin embedding, the skin tissue was cut into several continuous thin sections, each 4–6 µm thick. The first set of slides was stained with H&E, and five random fields were photographed at 200× magnification. Epidermal thickness was measured six times per field. The second set of sections was rehydrated, and antigen retrieval was performed using a pH 6 recovery solution. Immunohistochemistry (IHC) staining was conducted using rabbit anti-mouse myeloperoxidase antibody (Ab65871, Abcam Ltd, USA) at a concentration of 1 µg/mL, followed by goat anti-rabbit secondary antibody (GTX83399, GeneTEX, Taiwan) to stain antigen-positive cells in the dermis. Hematoxylin stain was then used as a counterstain. Six dermal areas were observed at 400× magnification. Based on the presence of positive multinucleated cells, infiltrating cell counts were categorized into five grades according to ISO 10993-6 2016: Table E1 [26]: 0 points (negative); 1 (1–5 positive cells per field); 2 (6–10 positive cells per field); 3 (abundant infiltration of positive cells); 4 (field densely packed with positive cells).

### Statistical analysis

Results are presented as the mean of the values, with error bars representing the 95% confidence intervals. The analysis package in GraphPad Prism 6.01 (Boston, MA) was used for statistical analysis. Statistical results were calculated by one-way ANOVA with Dunnett’s multiple comparisons test.

For statistical analysis of tissue histopathology, Fisher’s exact test was applied using the control group as the reference. A threshold of grade 3 or above was set for classification. The odds ratio (O.R.) was calculated using a one-tailed test. If the O.R. was less than 1, it indicated an effective treatment outcome; if greater than 1, the treatment was considered ineffective.

### Ethical statements

All animal works were supervised by a licensed veterinarian and performed in a GLP-compliant animal testing facility. All animal studies were reviewed and approved by the Institutional Animal Care and Use Committee (IACUC) of a qualified institute in Taiwan. All procedures involving study animals were conducted in a manner that avoided or minimized discomfort, distress or pain to the animals. Clinical symptoms were observed once per day on behavior, respiration, and appearance.

## Results

### Isolation and characterization of *C. asiatica* EVs

*C. asiatica* EVs were isolated from tissue culture *C. asiatica* by a centrifugation and filtration. The size distribution of EVs harvested was analyzed by NTA and showed that the particle has a mean size of 150 nm (Figure 1A). The morphological features shown by the TEM images can be observed as vesicles of approximately 100-150 nm in size with a lipid bilayer membrane structure (Figure 1B).

**Fig. 1.**
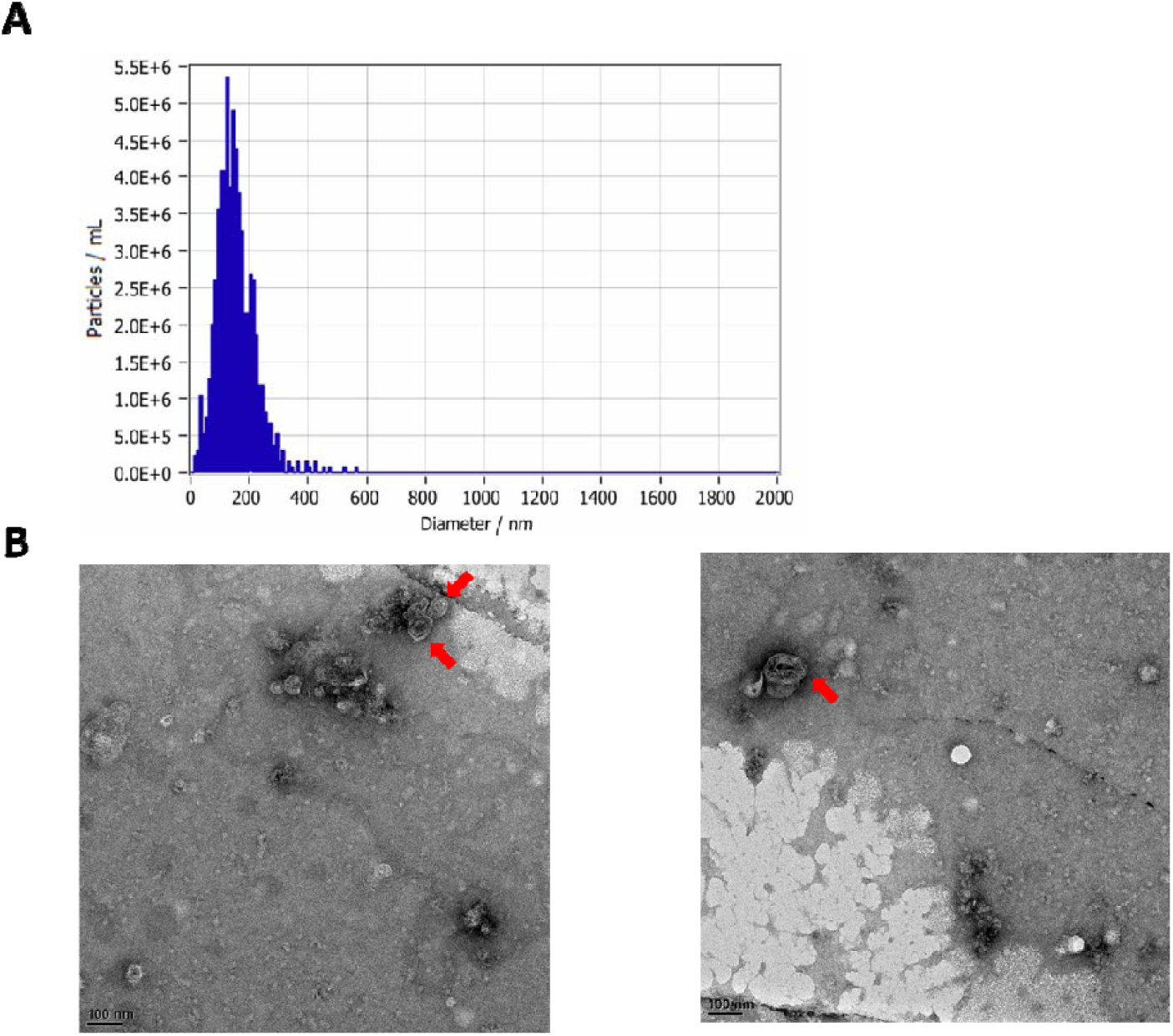
Size and morphology characterization of *C. asiatica*-derived EVs. A. Nanoparticle tracking analysis (NTA) of size and concentration distribution of EVs by using the ZetaView® system. B. Representative TEM images of purified *C. asiatica* EVs with red arrows showing vesicles with lipid bilayer membrane structure. Scale bar: 100 nm

### Cytotoxicity assessment

To investigate the effect of *C. asiatica* EVs in cells, an MTT assay was used to evaluate the effect of cell viability. B16F10 melanoma cells were treated with various concentrations of *C. asiatica* EVs and MTT assay was performed. The results showed no significant differences compared to the control group in this test (Figure 2). Thus, high concentrations of up to 5×10^9^ particles/mL of *C. asiatica* EVs do not exhibit cytotoxicity in B16F10 cells. In addition, no discernable differences in cell viability were detected in HaCaT and Detroit 551 cell lines, confirming that the EVs do not exhibit cytotoxicity in mouse or human cells (Figure S1).

**Fig. 2.**
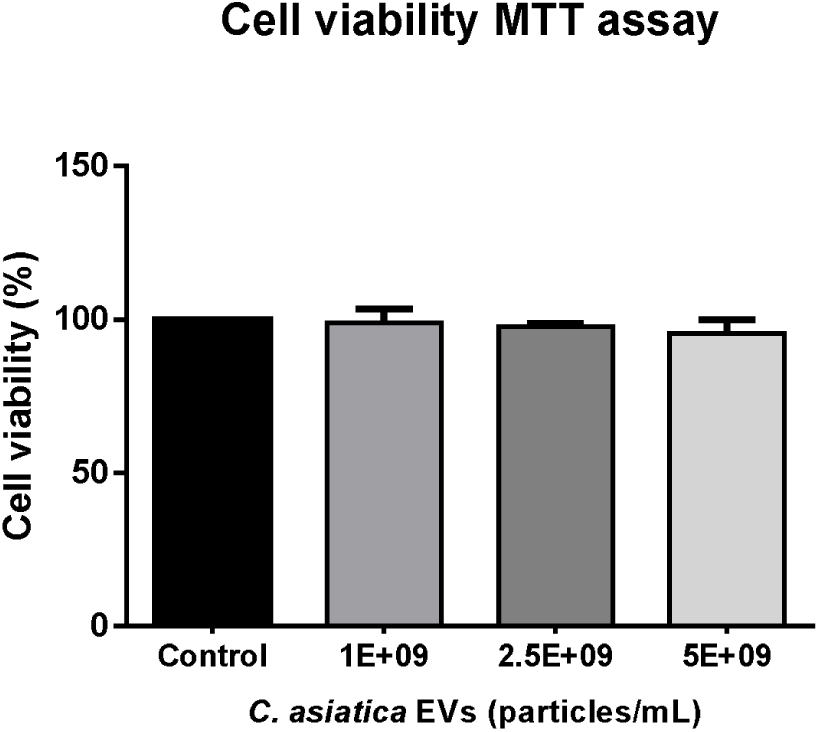
Cell viability assay. B16F10 cells were treated with different concentrations of *C. asiatica* EVs. After 24h treatment, cell viability was measured by the MTT assay method. Data are presented as mean ± SD (n=3). Statistical results were calculated by one-way ANOVA with Dunnett’s multiple comparisons test compared to the control group.

### Antioxidant activity assessment

A variety of naturally-occurring polyphenol compounds from *C. asiatica* extracts have been studied, which contribute to its antioxidant properties [5]. Here we investigated the total polyphenol content of *C. asiatica* EVs using an established method with the Folin-Ciocalteu reagent. The results showed that the gallic acid equivalent of *C. asiatica* EVs increased with the particle number, indicating that the antioxidant activity of the vesicles may be dose-dependent (Figure S2).

We also assessed the antioxidant activity of *C. asiatica* EVs by using two cell-free assays: ABTS and K_3_[Fe(CN)_6_] assays. At the higher concentrations, *C. asiatica* EVs exhibited comparable ABTS^+.^ scavenging capacity to positive controls vitamin C and BHA (Figure S3). For the K_3_[Fe(CN)_6_] metal chelating assay, the results showed that reducing capacities of EVs were also dose-dependent (Figure S4).

To further confirm the antioxidant capacity of *C. asiatica* EVs in B16F10 cells, intracellular reactive oxygen species (ROS) levels were measured by ROS scavenging rate using H_2_O_2_ at 24 mM to induce oxidative stress. After treatment, the intracellular ROS clearance was 53.86 `±` 0.94%, 59.32 `±` 1.68% and 66.27 `±` 1.10% for 1×10^9^, 2.5×10^9^ and 5×10^9^ (particles/mL) of *C. asiatica* EVs, respectively (Figure 3). The results showed that all concentrations of *C. asiatica* EVs significantly increased ROS scavenging activity compared to the control group (P<0.0001), demonstrating effects similar to positive control Trolox.

**Fig. 3.**
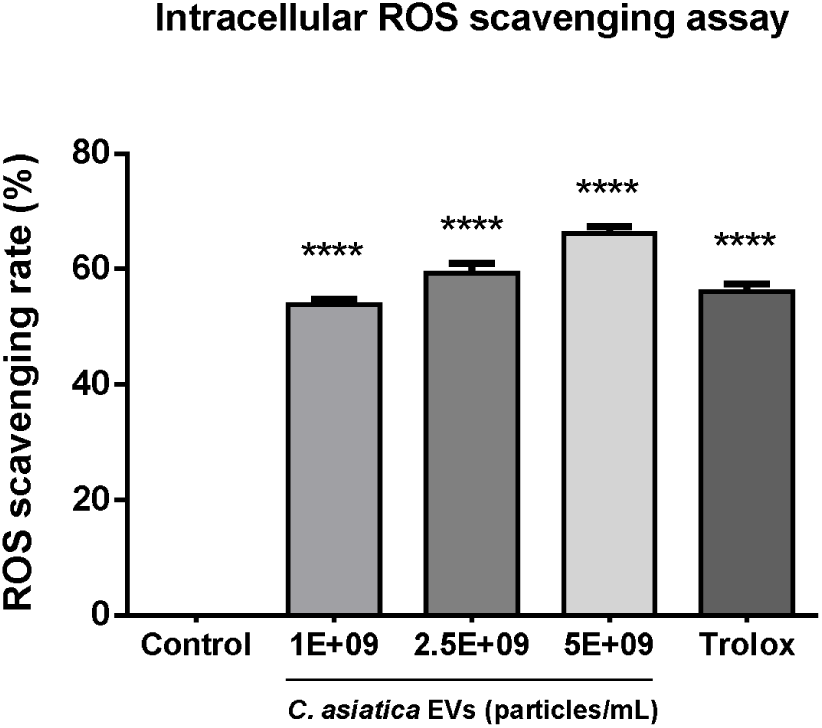
Intracellular ROS level. B16F10 cells were pretreated with various concentrations of *C. asiatica* EVs, Trolox (0.5 mg/mL, positive control), or left untreated as a control for 24 hours. The cells were then incubated with 24 mM H_2_O_2_ and DCFH-DA. The fluorescence intensities of DCF were measured to calculate the ROS levels. Data are presented as mean ± SD (n=3). Statistical analysis was calculated by one-way ANOVA with Dunnett’s multiple comparisons test, compared to the control group. ****P<0.0001

### Melanin inhibitory activity assessment

In addition to its antioxidant properties, *C. asiatica* has potential for skin whitening application through the inhibition of melanin synthesis [22]. First, we investigated its inhibition of melanin production in B16F10 melanoma cells treated with α-melanocyte-stimulating hormone (α-MSH). After *C. asiatica* EV treatment, the melanin content in the B16F10 cells was 93.55 ± 1.14%, 90.73 ± 1.17% and 83.69 ± 4.31% of control at 1×10^9^, 2.5×10^9^and 5×10^9^particles/mL of *C. asiatica* EVs, respectively (Figure 4A). For the positive standard arbutin (0.54 mg/mL), the remaining intracellular melanin content was 71.95 ± 2.44% of the control. Tyrosinase catalyzes the first two steps of melanin synthesis by hydroxylation of L-tyrosine to L-DOPA [23]. Therefore, to investigate the change in melanin synthesis via tyrosinase activity, we have carried out intracellular tyrosinase activity in B16F10 cells treated with α-MSH. After treatment, the remaining intracellular tyrosinase activity was 92.5 `±` 6.77%, 80.2 `±` 5.13% and 56.0 `±` 3.83% for the 1×10^9^, 2.5×10^9^ and 5×10^9^ particles/mL of either *C. asiatica* EVs, respectively (Figure 4B). In comparison, the remaining intracellular tyrosinase activity was 66 ± 10.53% of control after the cells were treated with arbutin, a known skin-whitening compound. The results indicated that 5×10^9^ particles/mL of *C. asiatica* EVs exhibited stronger inhibitory effect on α-MSH induced tyrosinase activity in B16F10 cells than arbutin, albeit with higher intracellular melanin content than arbutin.

**Fig. 4.**
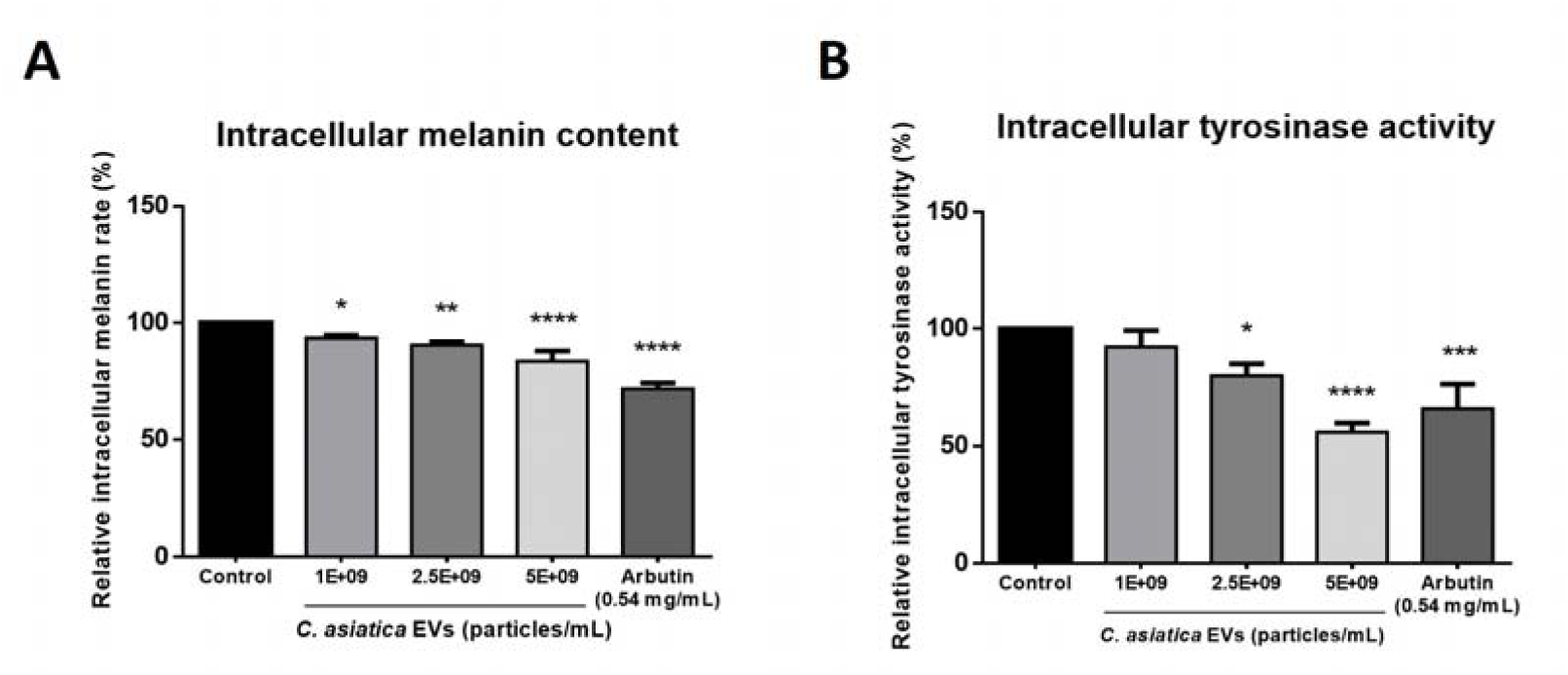
Effect of *C. asiatica* EVs on melanin production. Inhibitory effects of different concentrations of *C. asiatica* EVs on **A.** melanin content and **B.** intracellular tyrosinase activity in B16F10 cells. Data are presented as mean ± SD (n=3). Statistical results were calculated by one-way ANOVA with Dunnett’s multiple comparisons test compared to the control group. *P<0.05, **P<0.01, ***P<0.001, ****P<0.0001

### Effect of *C. asiatica* on skin integrity-related genes

One of the major components in skin tissue is type I collagen, which is derived from its precursor type I procollagen (PIP) secreted by skin fibroblasts, and lowered expression of PIP is associated with skin UV damage and senescence leading to wrinkle formation [24]. *C. asiatica* EVs increased production of PIP in Hs68 cells and induced a similar expression level of PIP to that of positive control TGF-β1 at the highest concentration tested (1×10^9^ particles/mL) (Figure S5). Aquaporin and filaggrin are important in the barrier function of skin by regulating water transport and water retention in skin [25]. Real-time PCR assay quantification of mRNA expression of aquaporin-3 (AQP3) and filaggrin (FLG) genes in HaCaT keratinocyte cells treated with *C. asiatica* EVs showed that *C. asiatica* EVs could upregulate the expression levels of AQP3 and FLG genes compared to the control group (Figure S6).

### Anti-inflammatory effects of *C. asiatica* EVs

UV exposure of keratinocytes can induce COX2 expression that increases pro-inflammatory prostalglandin synthesis, and *C. asiatica* triterpene madecassoside was found to inhibit UV-induced inflammation [22]. Here, we showed that the addition of *C. asiatica* EVs can lower the expression levels of COX2 in skin fibroblast Hs68 and keratinocyte HaCaT cells exposed to UV (Figure S7). Nitric oxide (NO) is an inflammatory response messenger secreted mainly by endothelial cells and macrophages, as well as keratinocytes, and exposure of skin to UV irradiation results in elevated levels of NO production [27]. We subjected *C. asiatica* EV-treated RAW 264.7 macrophage cultures to lipopolysaccharide (LPS), which is a TLR4 ligand and induces pro-inflammatory response [28]. The NO production was measured by the Griess reagent, which is converted to a pink color after reacting with nitrites. As shown in Figure 5, the amount of nitrite decreases in a dose-dependent manner in cells treated with increasing concentrations of EVs. Taken together, *C. asiatica* EVs were able to reduce UV-induced inflammatory response in a variety of cell types tested.

**Fig. 5.**
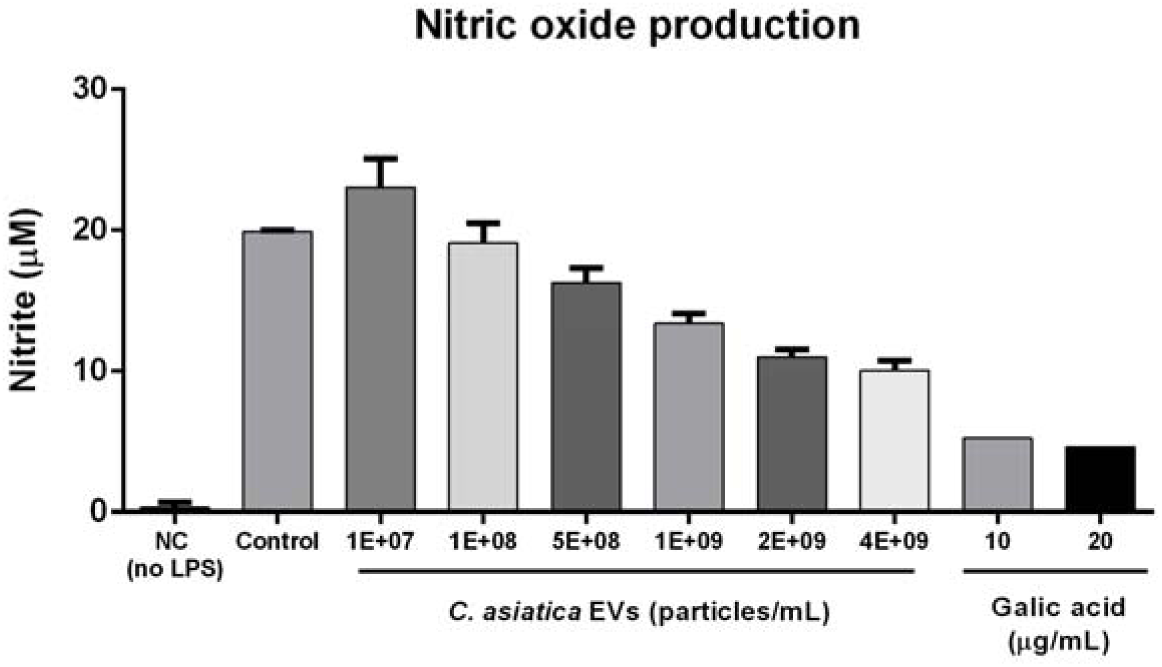
Nitric oxide (NO) production in RAW 264.7 macrophages. Cells were treated with different concentrations of *C. asiatica* EVs or gallic acid after induction by LPS. The resulting NO produced was detected by the Griess reagent.

### Effect of *C. asiatica* EV-based gel on UVB-induced skin damage in Mice

Next, we investigated whether a *C. asiatica* EV-based gel formulation (with or without the addition of *C. asiatica* extract TECA) could alleviate skin damage induced by UVB exposure. First, we evaluated the safety of the application of *C. asiatica* EV gel and we found no adverse effects in all groups that were applied with gel, regardless of the presence of *C. asiatica* EV or TECA (Table S1). There were also no significant weight differences between groups (p > 0.05), despite a slight decrease in body weight observed in all groups during the study (Table S2 and Figure S8).

In tissue histopathological analysis, the epidermis measurement results are summarized in Table 2 and Figure 6. The UVB-only group had slightly less epidermis thickness (126.66 ± 47.73 μm) compared to the control group (154.94 ± 58.14 μm), but the difference was not statistically significant. Treatment groups 1 and 2 had average epidermal thicknesses of 87.66 ± 35.32 μm and 86.09 ± 25.74 μm, respectively. Compared to the control group (154.94 ± 58.14 μm), the epidermal thicknesses were significantly (p < 0.01 for treatment 1 and p < 0.05 for treatment 2) reduced.

**Fig. 6.**
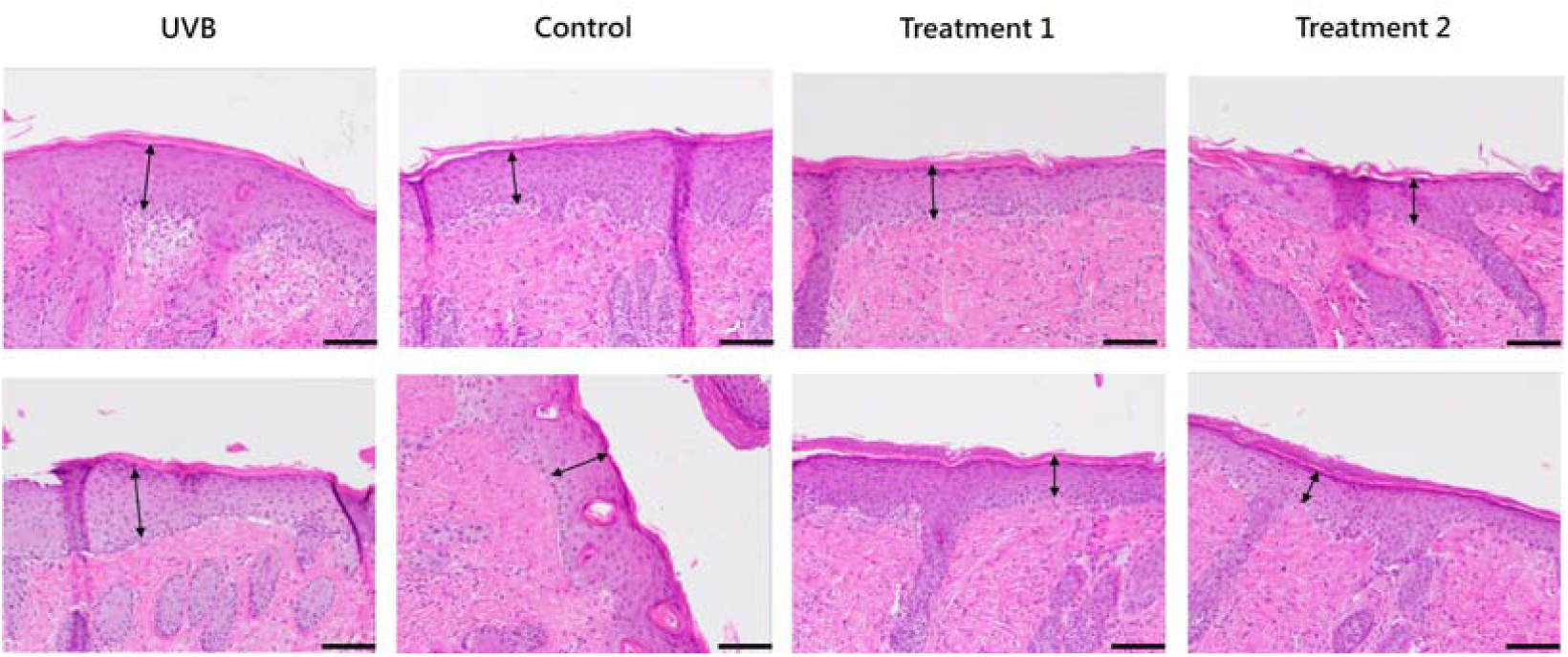
Representative examples of histopathological analysis of skin tissue sections on Day 7 of the study. Arrows indicate thickness of the epidermis. H&E stain, 200x magnification (scale bar = 100 µm)

**Table 2.**
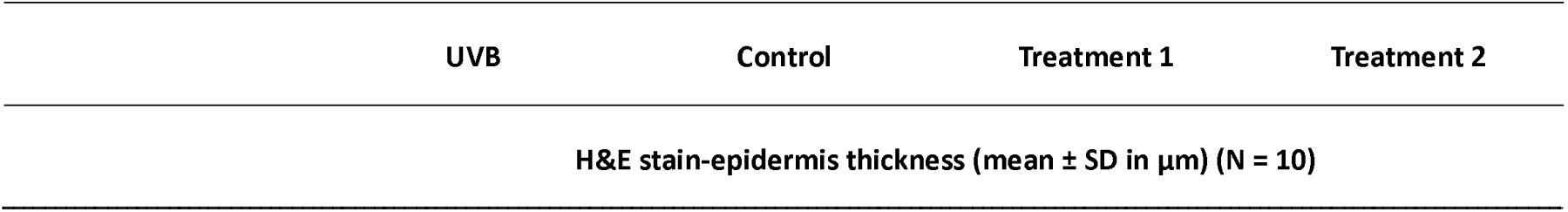
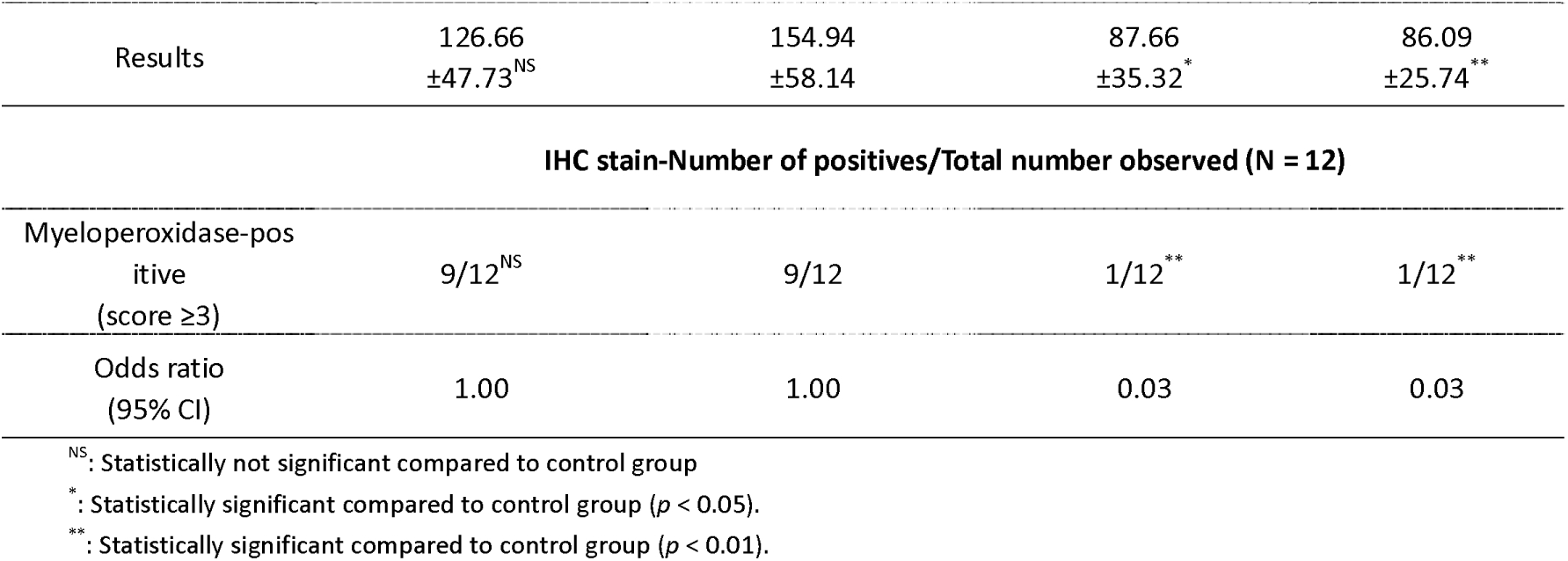
Tissue section measurement and observation. Results are taken from measurements of two skin sections (5 random fields for H&E stain, 6 random fields for IHC stain)

In addition, immunohistochemical staining of inflammation marker myeloperoxidase (MPO) was used to observe positive multinucleated cells in the dermis of UV-damaged areas (Table 2 and Figure 7). In both UVB-only and control groups, 75% of the observed fields exhibited extensive infiltration of positive multinucleated cells. The addition of *C. asiatica* EVs with or without TECA significantly reduced cellular infiltration (8.3% in both treatment groups). The odds ratios were also significantly lower in both treatment groups (0.03) compared to UVB and control groups (1.00) as calculated by Fisher’s exact test.

**Fig. 7.**
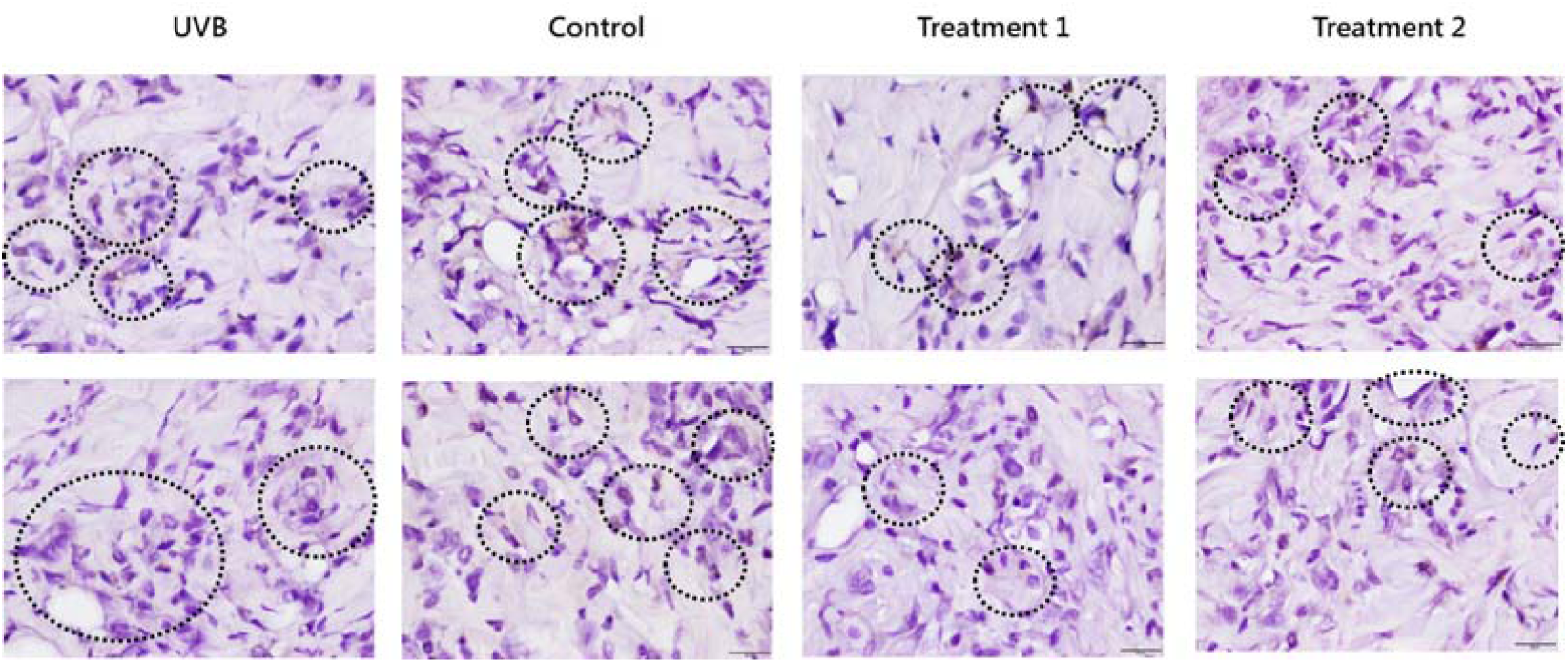
Representative examples of IHC staining of myeloperoxidase (in brown, indicated by circles) in skin tissue sections on Day 7 of the study. Mouse back skin sections were stained with rabbit anti-mouse myeloperoxidase antibody and then detected by HRP goat anti-rabbit antibody. 400x magnification (scale bar = 20 µm)

Gross appearance examination of mouse skin UVB exposure site on Day 6 of the study is shown in representative photographs in Figure 8. In UVB-only and blank gel control groups, inflammation and redness are still apparent at the UV wound site, with only a few newly repaired skin and abundant scarring. In contrast, for both treatment groups, the wound has been repaired and inflammation is absent at the site with minimal scarring.

**Fig. 8.**
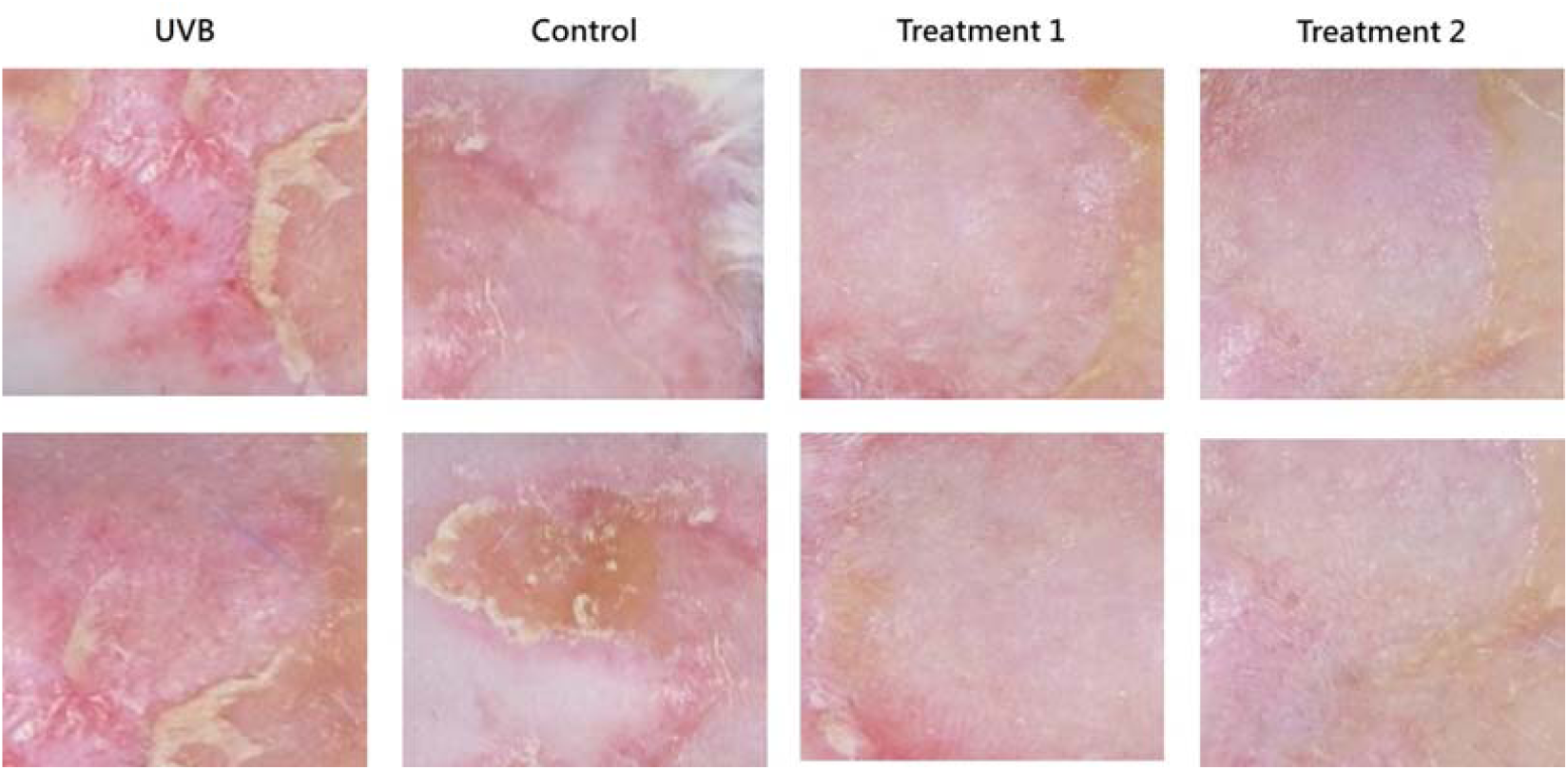
Representative examples of mouse skin appearance at the site of UVB exposure on Day 6 of the study.

## Discussion

In this study, we have isolated and characterized EVs secreted from *C. asiatica* tissue culture. The EVs have a size of approximately 150 nm and a lipid bilayer morphology similar to that previously reported for exosomes derived from *C. asiatica* and other plants [12, 13]. In a series of *in-vitro* experiments in B16F10 cell culture, we have demonstrated that the EVs are safe for cells and that EVs exhibited notable antioxidant, anti-melanogenic, and anti-inflammatory effects, suggesting potential applications in skin care and therapeutic treatments to combat oxidative stress and hyperpigmentation.

Another study also isolated exosome-like EVs from *C. asiatica* but with a smaller average particle size of 63.8 nm [14]. While the EVs from the above study exhibited cytotoxicity in HepG2 human cancer cells, we did not observe any effect on cell viability in mouse melanoma B16F10 cells or HaCaT and Detroit 551 human cell lines tested (Figures 2 and S1) [14]. The therapeutic properties of *C. asiatica* have mostly been attributed to the triterpenes and derivatives, including madecassoside, madecassic acid, asiaticoside, and asiatic acid, and these are among the most well-studied compounds in the plant [3–5]. In our study, we observed potent antioxidant effects from *C. asiatica*; however, further study is required to determine specifically how contents within EVs, such as phytochemicals, proteins, and nucleic acids, can contribute to the observed effects.

In addition to phytochemicals and proteins, small RNAs and miRNAs in EVs are important modulators of target effect proteins. RNA-binding proteins such as Argonaute 1, RNA helicases, and Annexins, may be involved in packing RNA into EVs [10, 29] While our study did not include direct profiling of miRNAs, a recent preprint study has provided a comprehensive analysis of the transcriptome human skin keratinocytes after treatment of *C. asiatica* exosomes which revealed 11 novel miRNAs packed within the exosomes [13]. These miRNAs can modulate genes related to skin integrity and inflammation, skin integrity and barrier-related genes and such genes were upregulated after treatment by *C. asiatica* EVs, which we have also observed in this study with increasing gene expression of PIP1, AQP3, and FLG (Figures S5 and S6) [13]. The above study also found that *C. asiatica* exosomes induced higher gene expression than *C. asiatica* extracts, possibly due to better uptake of exosomes by skin cells and the effects of miRNAs packaged within [13]. Thus, while *C. asiatica* extracts are rich in triterpenoids such as madecassoside. EVs offer an additional layer of bioactivity through the encapsulated biomolecules, such as small RNAs and proteins. Unlike plant extracts, EVs can facilitate targeted delivery for improved penetration and cross-kingdom gene regulation [8, 11].

In *C. asiatica*, madecassoside by itself has the ability to inhibit UV-induced melanin synthesis in a keratinocyte and melanocyte co-culture system as well as artificially tanned human skin [22]. Other plant-derived EVs also have anti-melanogenic effect. For example, *Dendropanax morbifera* EVs can inhibit the expression of melanogenesis genes such as MITF, TYR, TRP-1, and TRP-2 [9, 28]. In our study, the EVs appeared to modulate melanogenesis pathways in cell culture, likely through the inhibition of tyrosinase activity, which is a key enzyme in melanin synthesis [23]. The mechanism of anti-melanogenesis by *C. asiatica* includes inhibition of inflammation, as observed in previous studies [24]. and in our study, where we have observed that *C. asiatica* can reduce pro-inflammatory gene expression, such as COX2 and decrease the production of NO (Figures 5 and S7). As melanin overproduction contributes to skin conditions like freckles and nevus formation, and age spot development, plant-derived EVs with anti-melanogenic properties represent an economical and natural alternative to conventional treatments [29].

In the animal study, mice were exposed to UVB at 120 mJ/cm² for 5 minutes to induce photoaging and skin damage. This was followed by daily application of EV gel for seven consecutive days. Photoaging leads to degeneration of elastic fibers and collagen loss, resulting in epidermal thickening of the stratum corneum [30]. In both treatment groups consist of *C. asiatica* EV gels with or without TECA, we observed reduced epidermal thickening in skin sections and reduced cell infiltration due to inflammation characterized using IHC staining, suggesting a potential skin repair effect. However, in the control group with blank gel application, a thicker, albeit not statistically significant, epidermis layer was observed than UVB-only group, possibly due to mechanical irritation from the gel application process. These preliminary results suggest that EVs may help to alleviate UV-induced inflammation but are unable to reverse UV-induced epidermal damage.

We have recently conducted a 28⍰day clinical trial to evaluate the efficacy of a topical *C.⍰asiatica* EV–based formulation in 20 healthy adults [31]. Application of the topical product twice daily resulted in statistically significant improvements in skin hydration, elasticity, complexion brightness, wrinkle reduction, redness, pore size, and reduced melanin content over the study period [31]. This clinical data complements our *in vitro* and murine model findings by demonstrating that the skin⍰protective activities we observed, especially in reducing melanin, inflammation, and improving barrier integrity, translate into measurable cosmetic and functional benefits in humans. In particular, *C. asiatica* EVs upregulated AQP3, FLG, and procollagen genes *in vitro*, correlating with the increased hydration and elasticity in clinical observations, as well as significant decreases in melanin content in trial participants, which also support our *in vitro* findings of tyrosinase inhibition and melanin suppression. The reduction in redness during the clinical study period is reflected in the anti-inflammatory pathways we demonstrated here, including COX2 downregulation *in vitro* and MPO reduction *in vivo*. The tissue culture micropropagation technology for *C. asiatica* has been established as a sustainable source of *C. asiatica* [32]. This ensures its conservation in nature while providing a constant yield of raw material, akin to the use of bioreactors in biopharmaceutical drug and vaccine production. Using plant tissue culture as a source for EV production presents several advantages over traditional methods of harvesting from wild plants. First, tissue culture allows for consistent, scalable production of EVs with genetic uniformity, bypassing the variability associated with environmental factors that affect wild plants, such as seasonal changes, soil quality, and climate/geographical conditions [15, 29]. This standardization is particularly important for achieving reproducible results in therapeutic and cosmetic applications, where consistent quality and efficacy are crucial in ensuring the safety and efficacy of the product. Additionally, tissue culture can be optimized to enhance the yield of specific bioactive compounds through controlled environmental conditions and the use of elicitors [15].

This study has several limitations. First, although we demonstrated the biological activity of *C. asiatica* EVs through various *in vitro* assays and a short-term UVB-induced mouse model, the precise molecular mechanisms, such as the role of small RNAs or miRNAs within the EVs, were not directly investigated. Second, the skin penetration and stability of the EVs in the topical gel formulation have not been fully evaluated. Third, the animal study involved a relatively small sample size (n=3 per group) and a limited observation period of 7 days, which may not fully capture the long-term and broader effects of EVs on skin quality and inflammatory responses. Future research should include transcriptomic or proteomic profiling of EV content and compare it to that of a previous study on *C. asiatica* EV transcriptome analysis [13].

In conclusion, this *in vitro* and animal model study demonstrated the antioxidant, anti-melanogenic and anti-inflammatory potential of *C. asiatica*-derived EVs from tissue culture. The advantages of tissue culture over wild plants in terms of sustainability, consistency, and the ability to enhance bioactive compound production further support the use of this approach in developing plant-based EV therapies. Future studies could focus on characterizing bioactive molecules and miRNAs within these EVs, as well as exploring their safety and efficacy in *in-vivo* models. Ultimately, this work paves the way for the sustainable production and utilization of plant-derived EVs in skin care cosmetics and wound healing applications.

## Acknowledgements

We thank the team at Schweitzer Biotech Company for reviewing and providing feedback for the manuscript preparation.

