## Supplement Figures for "*In Vitro* Characterization of Extracellular Vesicles from the Medicinal Plant *Centella asiatica* for Aesthetic Applications"

**Supplemental Figures**

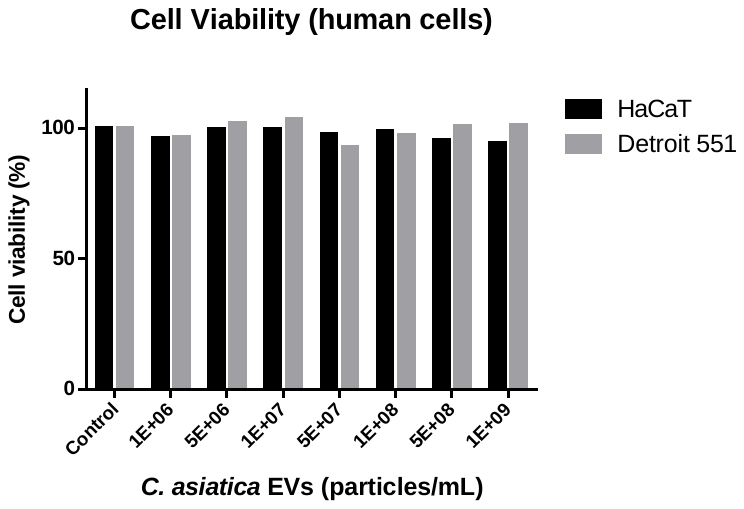

**Fig. S1** Cell viability assay in human cells. Human cell lines HaCaT and Detroit 551 were treated with different concentrations of *C. asiatica* EVs. After 24h of treatment, cell viability was measured by the alamarBlue® reagent method.

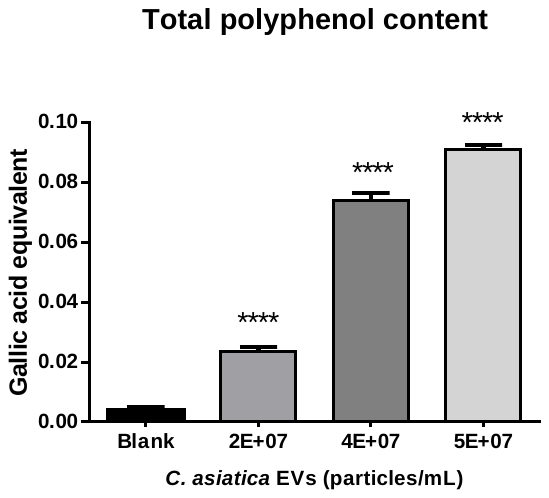

**Fig. S2** Total polyphenol content. The total polyphenol content of *C. asiatica* EVs using the Folin-Ciocalteu method. Data are presented as mean ± SD (n=3). Statistical results were calculated by one-way ANOVA with Dunnett’s multiple comparisons test compared to the control group. ****P<0.0001

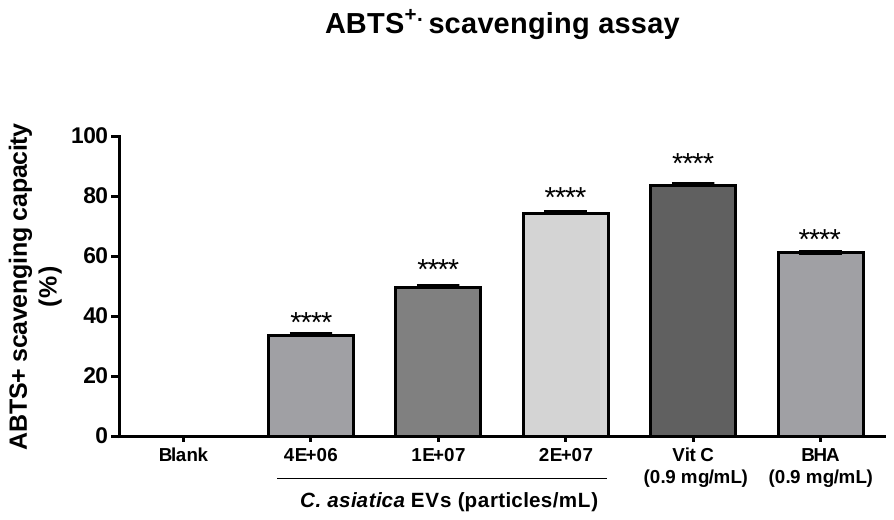

**Fig. S3** ABTS free radical scavenging activity. Free radical scavenging activities of *C. asiatica* EVs and vitamin C and BHA (as positive controls) were measured by ABTS free radical assay. Data are presented as mean ± SD (n=3). Statistical results were calculated by one-way ANOVA with Dunnett’s multiple comparisons test compared to the blank group.

****P<0.0001

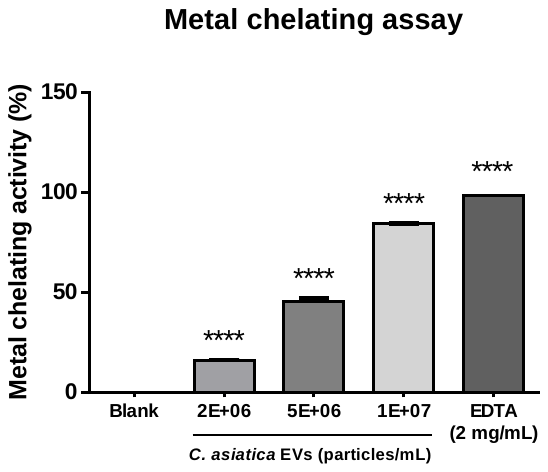

**Fig. S4** Metal chelating activity. The metal chelating activity of different concentrations of *C. asiatica* EVs and EDTA (2 mg/mL, positive control). Data are presented as mean ± SD (n=3). Statistical results were calculated by one-way ANOVA with Dunnett’s multiple comparisons test compared to the control group. ****P<0.0001

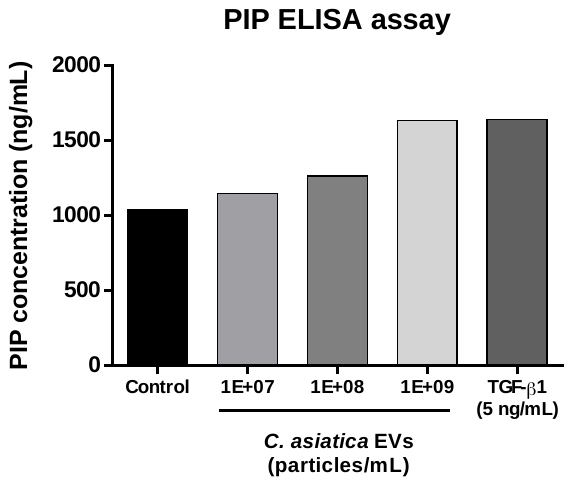

**Fig. S5** Concentration of secreted procollagen type I. The procollagen type I level of different

concentrations of *C. asiatica* EVs in Hs68 cells were measured using ELISA and quantified by a standard curve.

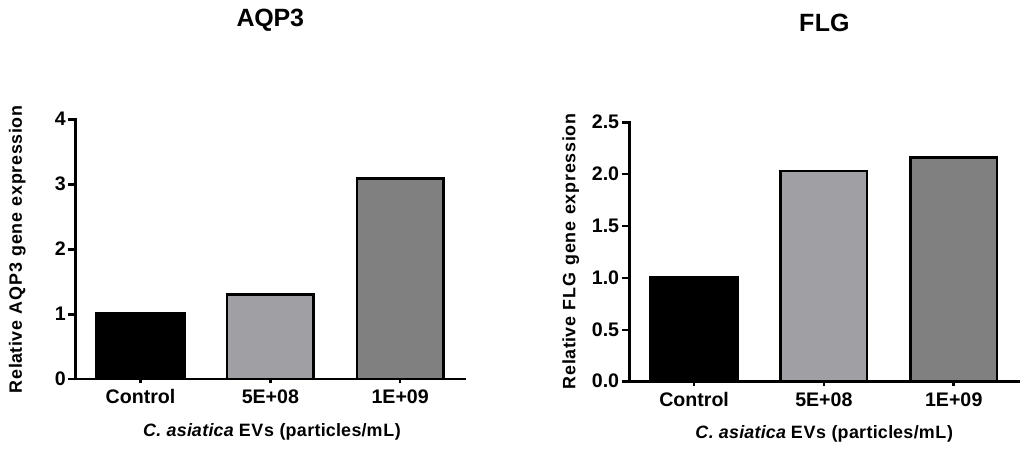

**Fig. S6** Expression of moisturizing and skin barrier improvement-related genes in different concentrations of *C. asiatica* EV-treated HaCaT cells. The results are expressed as the mRNA expression level of AQP3 and FLG relative to GAPDH expression.

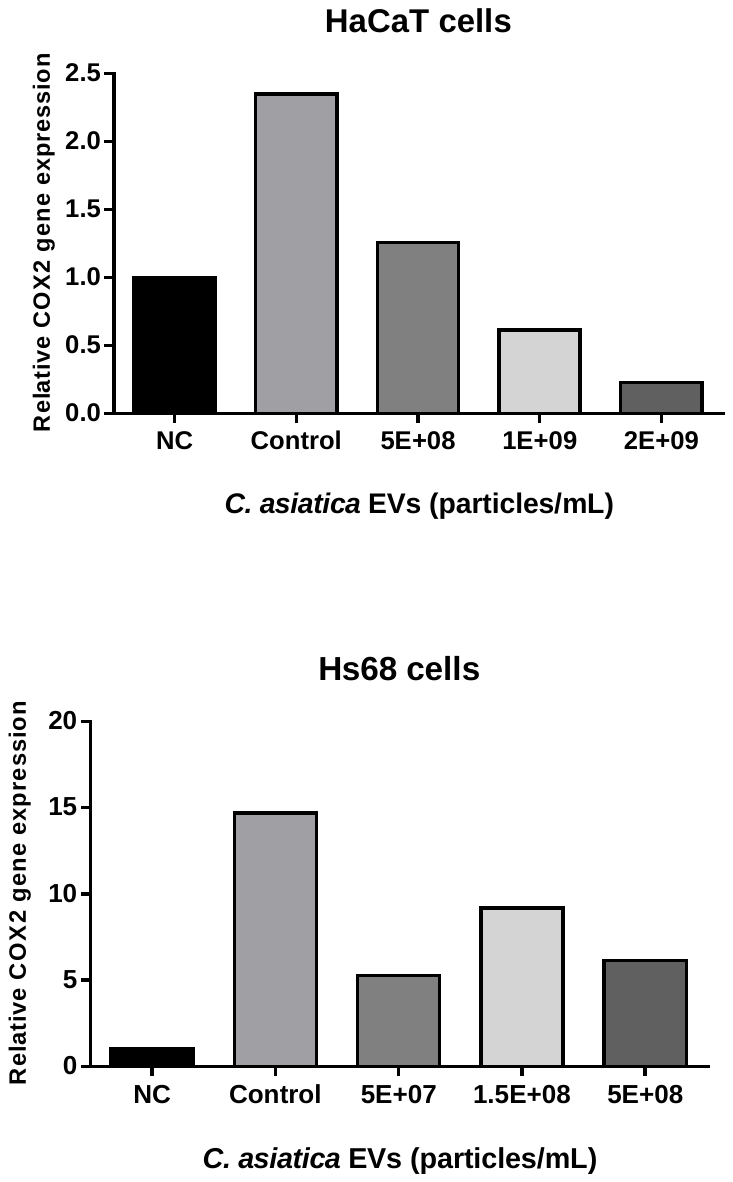

**Fig. S7** Expression of pro-inflammatory genes in cells exposed to a 90 mJ dose of UVB after treatment by different concentrations of *C. asiatica* EVs. The results are expressed as the mRNA expression level of COX2 relative to GAPDH expression.

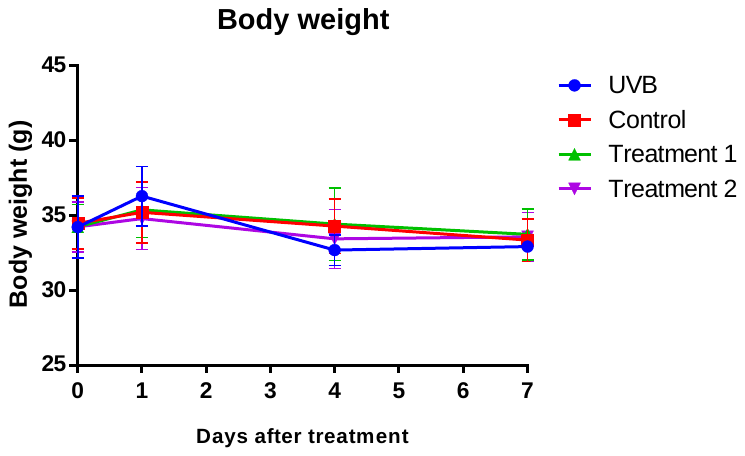

**Fig. S8** Body weight (in grams) of mice exposed to UVB on Day 0 and treated daily with blank gel (control), *C. asiatica* EV gel (treatment 1), or *C. asiatica* EV gel plus *C. asiatica* extract TECA (treatment 2). Results are presented as the mean body weight of each group of mice taken on Days 0, 1, 4, and 7 after UVB exposure, with error bars as the standard deviation of the mean body weight.
