## Supplement Tables for "*In Vitro* Characterization of Extracellular Vesicles from the Medicinal Plant *Centella asiatica* for Aesthetic Applications"

**Table S1** Clinical observation of animals ^(1)^

| Groups | Animal exhibiting abnormality | | | | | | | Total occurrence rate (N/N) ^(2)^ |
| --- | --- | --- | --- | --- | --- | --- | --- | --- |
|  | Day 1 | Day 2 | Day 3 | Day 4 | Day 5 | Day 6 | Day 7 |  |
| UVB | 0 | 0 | 0 | 0 | 0 | 0 | 0 | 0/3 |
| Control | 0 | 0 | 0 | 0 | 0 | 0 | 0 | 0/3 |
| Treatment 1 | 0 | 0 | 0 | 0 | 0 | 0 | 0 | 0/3 |
| Treatment 2 | 0 | 0 | 0 | 0 | 0 | 0 | 0 | 0/3 |

^(1)^Clinical observation items： Appearance, mood, behavior, respiration, mouth and nose, eyes, skin, digestion, or metabolism

^(2)^ N/N： Number of abnormal animals/Total number of animals observed

**Table S2** Body weight of animals over the course of the study (in grams).

| **Groups** | **Animal** | **Day 0** | **Day 1** | **Day 4** | **Day 7** |
| --- | --- | --- | --- | --- | --- |
| **UVB** | B1 | 36.51 | 38.56 | 33.82 | 33.15 |
|  | B2 | 33.62 | 35.22 | 31.85 | 32.91 |
|  | B3 | 32.50 | 35.04 | 32.36 | 32.68 |
|  | **Mean** | **34.21** | **36.27** | **32.68** | **32.91** |
|  | **SD** | **2.07** | **1.98** | **1.02** | **0.23** |
| **Control** | R1 | 33.88 | 34.44 | 34.08 | 33.55 |
|  | O2 | 33.12 | 33.64 | 32.57 | 31.82 |
|  | R3 | 36.37 | 37.48 | 36.18 | 34.65 |
|  | **Mean** | **34.46** | **35.19** | **34.28** | **33.34** |
|  | **SD** | **1.70** | **2.03** | **1.81** | **1.43** |
| **Treatment 1** | **O1** | 34.93 | 35.84 | 35.57 | 34.54 |
|  | **O3** | 32.52 | 33.31 | 31.63 | 31.76 |
|  | **O4** | 35.22 | 36.90 | 36.03 | 34.85 |
|  | **Mean** | **34.22** | **35.35** | **34.41** | **33.72** |
|  | **SD** | **1.48** | **1.84** | **2.42** | **1.70** |
| **Treatment 2** | **G1** | 35.90 | 36.51 | 35.18 | 35.09 |
|  | **G2** | 34.22 | 35.32 | 33.77 | 33.73 |
|  | **G4** | 32.56 | 32.51 | 31.31 | 31.85 |
|  | **Mean** | **34.22** | **34.78** | **33.42** | **33.56** |
|  | **SD** | **1.67** | **2.05** | **1.96** | **1.63** |
